## supplementary material for "Mutation-induced infections of phage-plasmids"

The PDF file includes

**Supplementary tables S1**

**Supplementary figures S1 to S12**

**Table S1 Details of all 21 mutations identified in 15 independent lines of populations.** These include 9 lines (populations a-i) of temporally-tracked populations with genomic sequencing, 3 lines with transcriptomic sequencing (populations j-l) and 3 lines of populations that we only sequenced colonies at the end timepoint (populations m-o). Notably, some mutations happened in parallel in two independent lines, which are marked in red.

| | position | population | source | reference | alteration |
|---|---|---|---|---|---|
| 1 | 11211 | j | transcriptomics | GTTTTTT | GTTTTTTT |
| 2 | 11345 | m | genomics | T | G |
| 3 | 11345 | e | genomics | T | G |
| 4 | 11363 | g | genomics | AG | AGG |
| 5 | 11364 | h | genomics | GT | GTT |
| 6 | 11366 | n | genomics | A | AT |
| 7 | 11366 | c | genomics | A | T |
| 8 | 11732 | b | genomics | TGGTGAGGT | TGGTGAGGTGAGGT |
| 9 | 11736 | i | genomics | GA | G |
| 10 | 11750 | b | genomics | CAA | CAAA |
| 11 | 11750 | d | genomics | CAA | CAAAA |
| 12 | 11771 | d | genomics | CG | CGG |
| 13 | 11806 | l | transcriptomics | TC | T |
| 14 | 11833 | k | transcriptomics | G | GC |
| 15 | 11840 | a | genomics | ATTT | ATT |
| 16 | 11853 | i | genomics | C | T |
| 17 | 11887 | a | genomics | CG | C |
| 18 | 11897 | o | genomics | GAAAAA | GAAAA |
| 19 | 11897 | b | genomics | GAAAAA | GAAAA |
| 20 | 11942 | a | genomics | CTT | CT |
| 21 | 11942 | f | genomics | CTT | CT |

**Figure S1 Assembly graph of *Tritonibacter mobilis* A3R06 complete genome.** The whole genome has 4.65 million base pairs (Mbp), with a chromosome of 3.2 Mbp plus four (mega)plasmids of 1.2 Mbp, 0.1 Mbp, 78 thousand base pairs (Kbp) and 42 (Kbp).

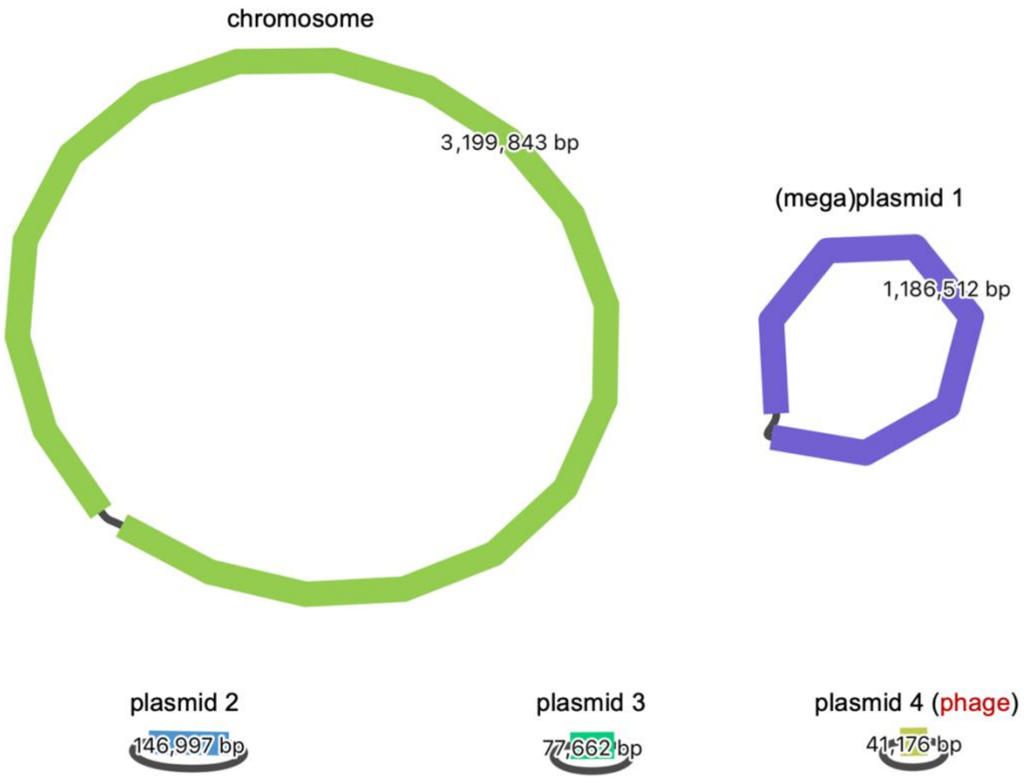

**Figure S2 Genome map of *Tritonibacter mobilis* A3R06 phage-plasmid.** The phage-plasmid has 51 predicted genes, including phage structural genes (e.g., a phage head, tail, capsid and portal proteins, purple), lysozyme (orange), cell wall hydrolases (orange) as well as a C1-type phage repressor (red). The rest of the genes are involved in plasmid stability, replication and segregation, such as the y*oeB*-*yefM* toxin-antitoxin system (pink), the *parAB* plasmid segregation system (blue), P4-family plasmid primase (yellow) as well as various methylases (green) for epigenetic modifications.

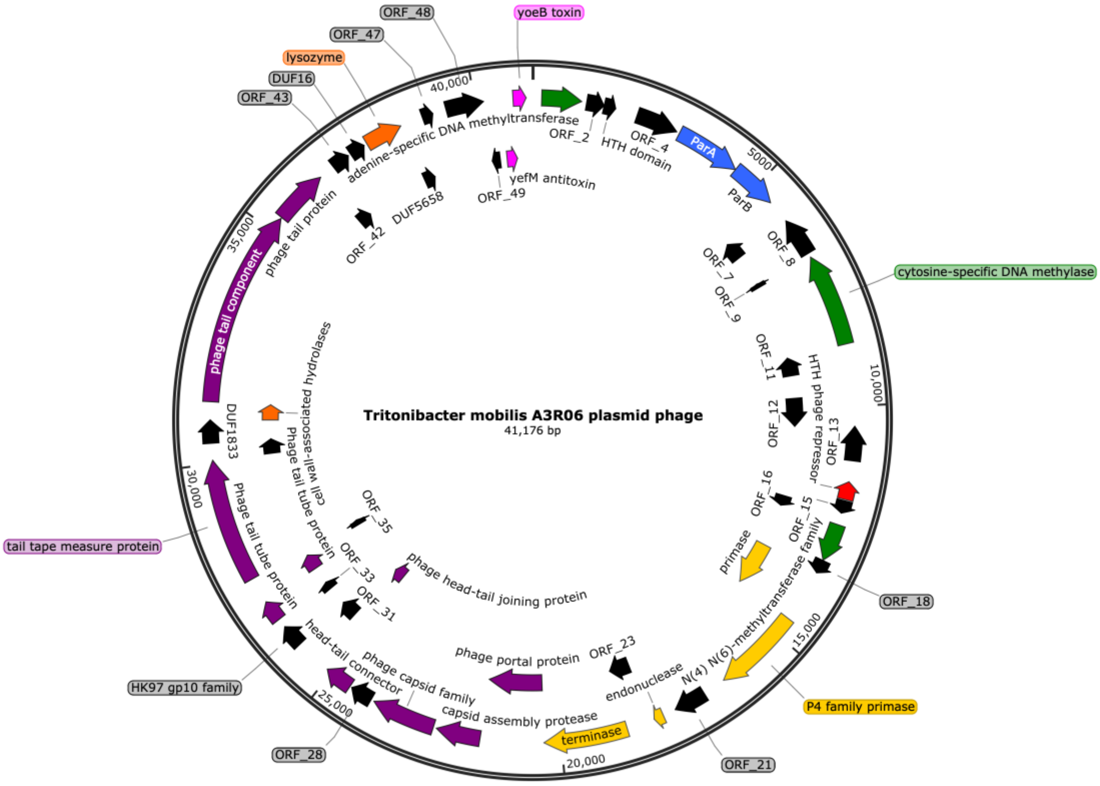

**Figure S3 Manually measured growth curve of *Tritonibacter mobilis* A3R06.** Culture growth followed an experimental physiology protocol including appropriate pre-culturing and temperature-controlled shaking conditions, as detailed in Methods. Guided by the measured doubling time of ~ 4h, we set the period of each serial dilution to be 24 hours and dilution factor to be 1/64. This translated to ~6 generations per cycle and the cells are always in the exponential phase throughout the cycles.

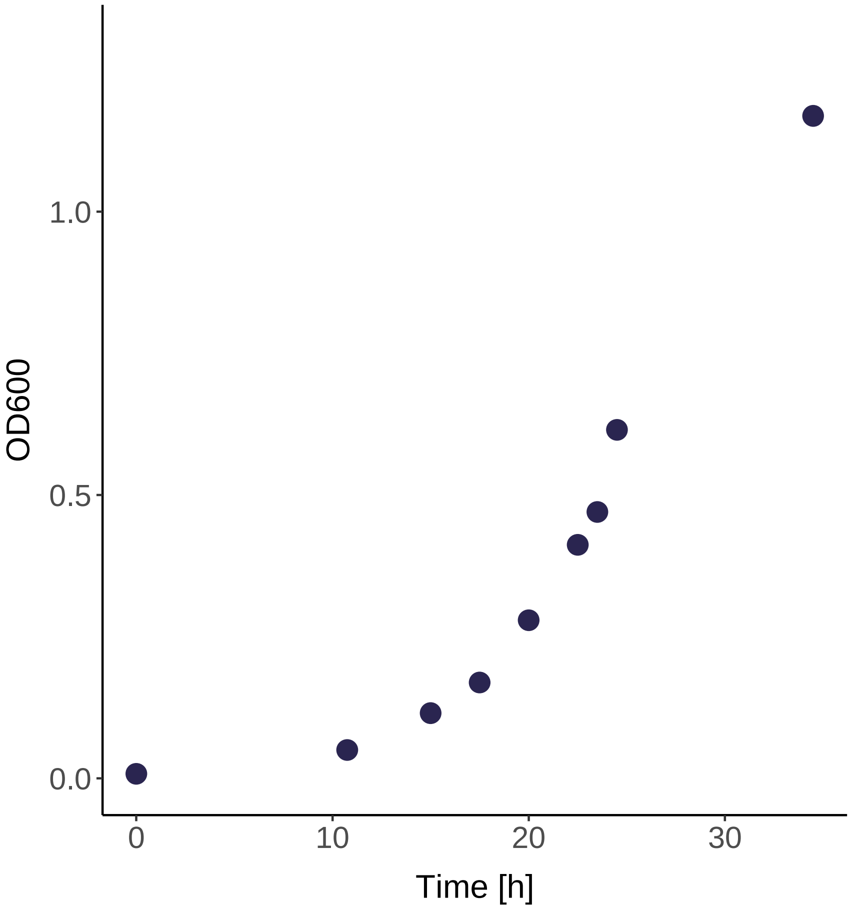

**Figure S4 Eco-evolutionary dynamics of the other 8 independent lines of populations that were temporally-tracked with genomic sequencing.** Each line of population (populations a-i) started from a different single colony. Bacterial culture was collected at the end of each dilution cycle, for which we did genome sequencing to determine the genotype frequencies of mutations (A) as well as the relative copy number of the phage-plasmid compared to the host chromosome (C). OD600 was also measured at the end of each dilution cycle (B).

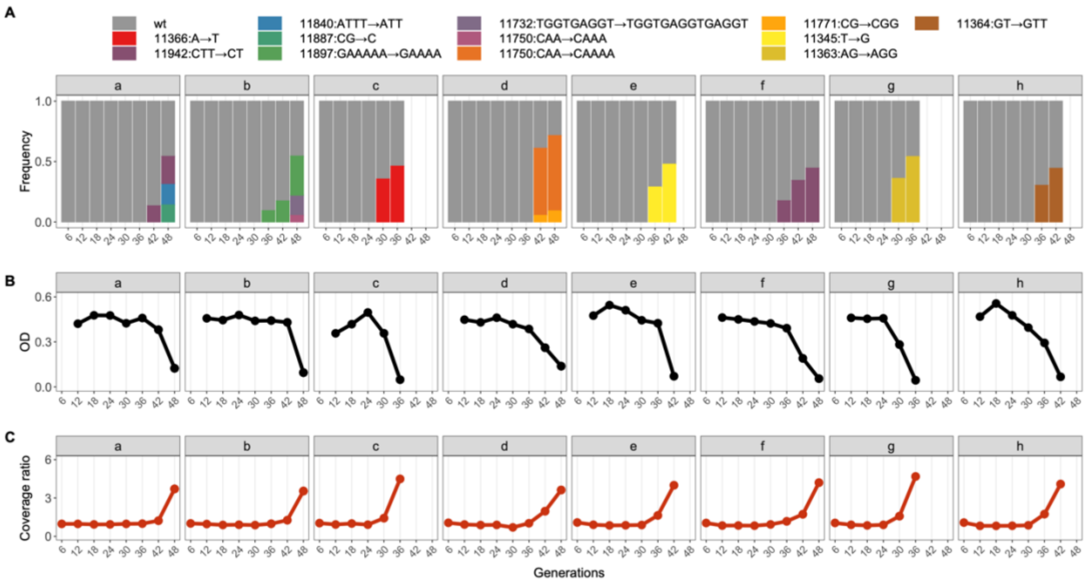

**Figure S5 Images of *Tritonibacter mobilis* A3R06 culture.** (A) Liquid culture was planktonic without phage-plasmid induction. (B) Culture became clumpy with cell aggregates when the mutated phage-plasmid was induced. (C) Fluorescent microscopic image of the cell aggregates. Green light by SYTO9 visualize live cells by binding to intracellular DNA while red light by propidium iodide visualize dead cells by binding to extracellular DNA.

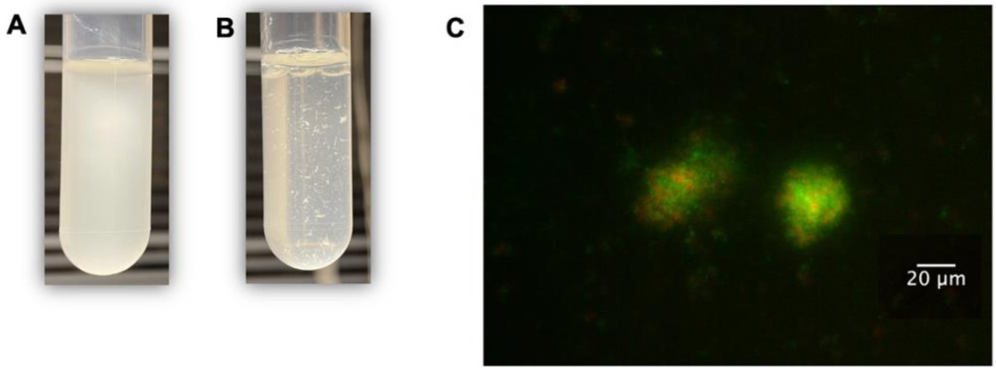

**Figure S6 Expression levels of phage-plasmid genes before and after the mutation was observed.** Genes related to plasmid replication and stability were constituently expressed even when the phage is lysogenic. Genes that are related to phage production became highly up-regulated after productive switch.

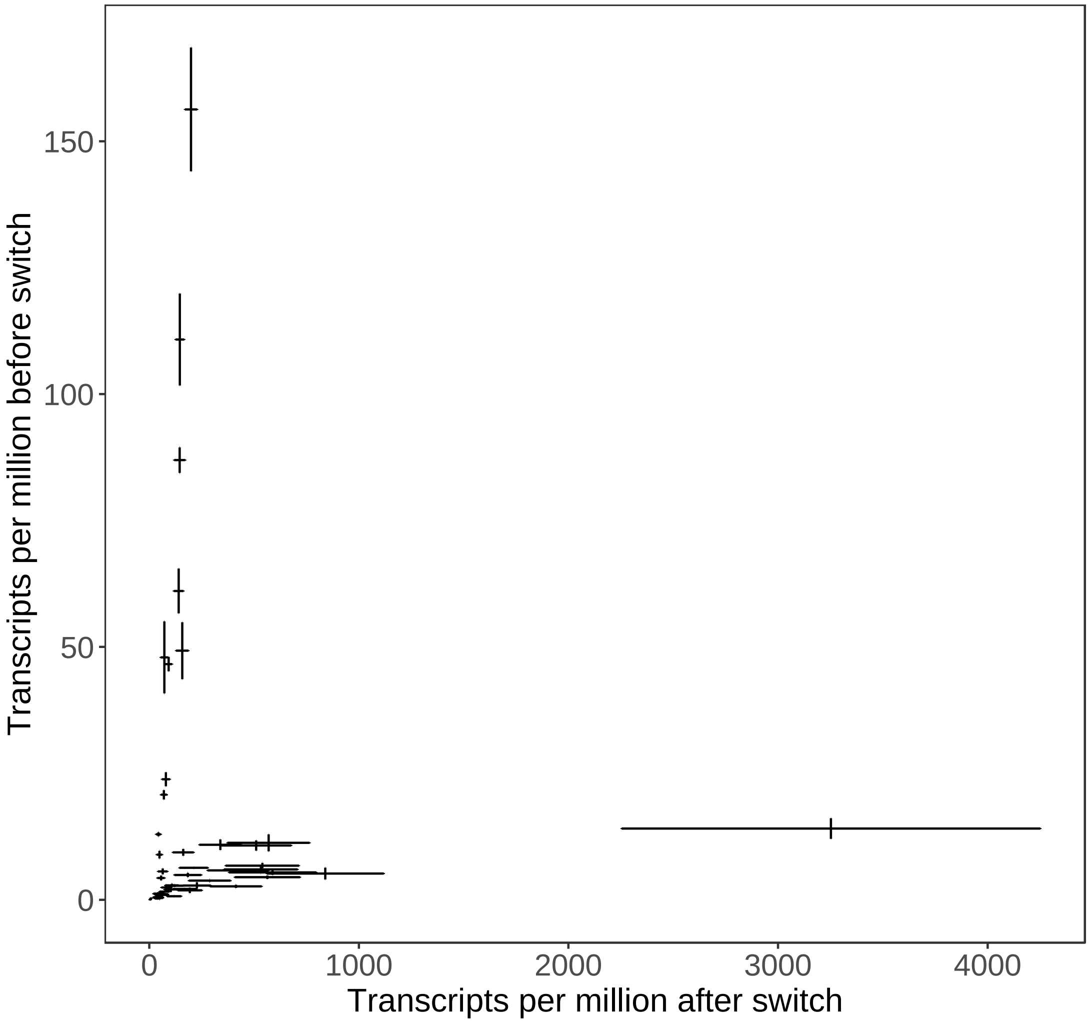

**Figure S7 Schematic illustration of segregational drift.** Here we showed a simplest scenario where two copies of phage-plasmids were randomly distributed into daughter cells with equal opportunity, while the copy number of phage-plasmids per cell remained constant. Following a Binomial distribution, we have 1/3 probability of generating two homozygotic descendants and 2/3 probability of generating two heterozygotic descendants.

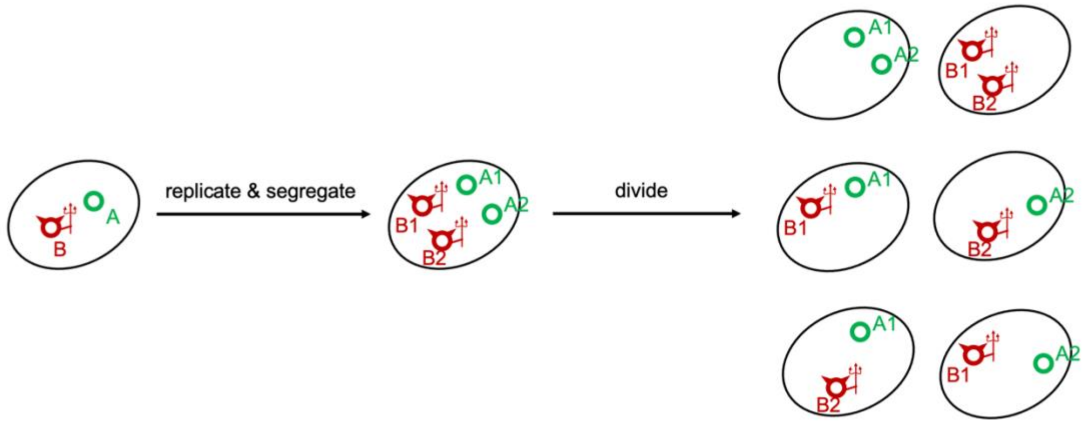

**Figure S8 Copy number of phage-plasmid per host cell.** Post segregational drift descendant populations carrying only wild-type phage-plasmid harbored a significantly elevated copy number of the phage-plasmid compared to the original host population before mutation-driven phage-plasmid induction. Notably, the average copy number in post segregational drift descendant populations was not an integer but between one and two, suggesting that some individuals may lose some copies of phage-plasmids during cell division.

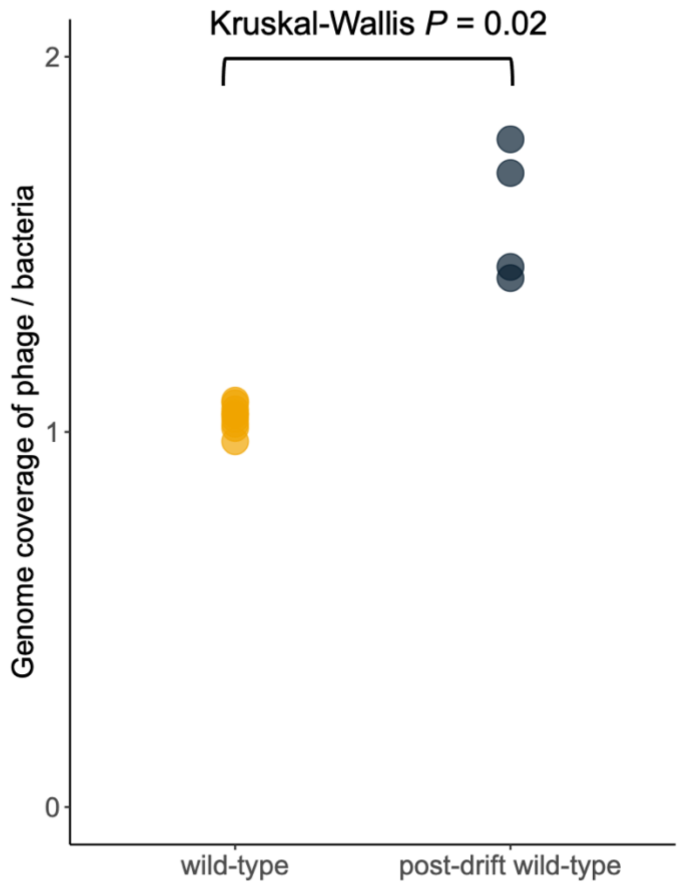

**Figure S9 Post segregational drift descendant populations carrying only wild-type phage-plasmid but with a higher copy number became more resistant to mutation-driven phage-plasmid induction.** Compared to the timing of 30~50 generations for populations carrying single copy of phage-plasmid, these post segregational drift populations were resistant to phage induction for 66 generations or longer under the same experimental serial dilution regime. Each color represents an independent line of post-segregational drift population under serial dilution scheme.

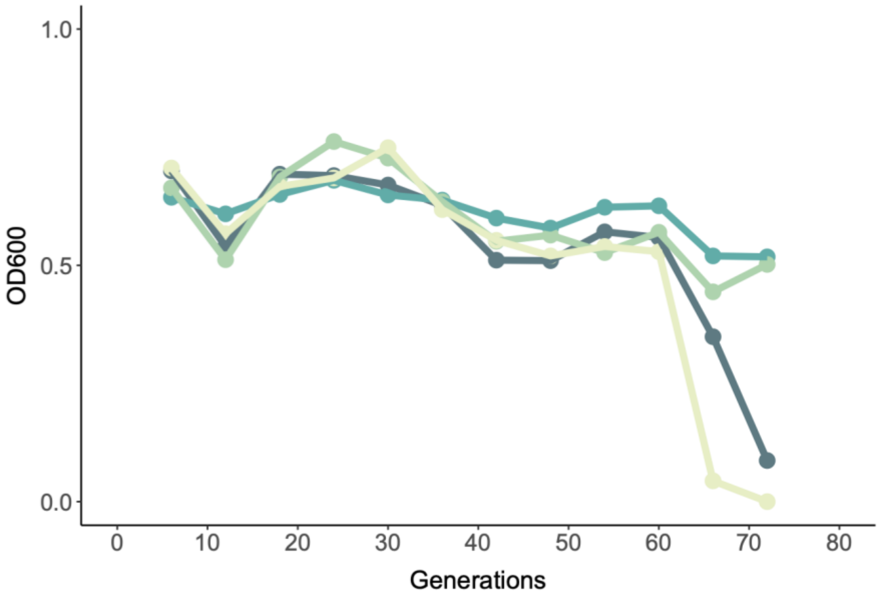

**Figure S10 The relationship between reinfection efficiency and saturating genotypic frequency in model simulation.** In our model, reinfection efficiency (R) indicates the average number of released mutated phages that successfully re-infect a host cell per host cell lysed. The resulting saturating frequency of the mutated genotype increases when reinfection efficiency increases, but the increase decelerates indicating the mutated genotype never reaches fixation. We found that $R = 5$ best fitted the experimentally observation of the saturating genotypic frequency of roughly 0.5.

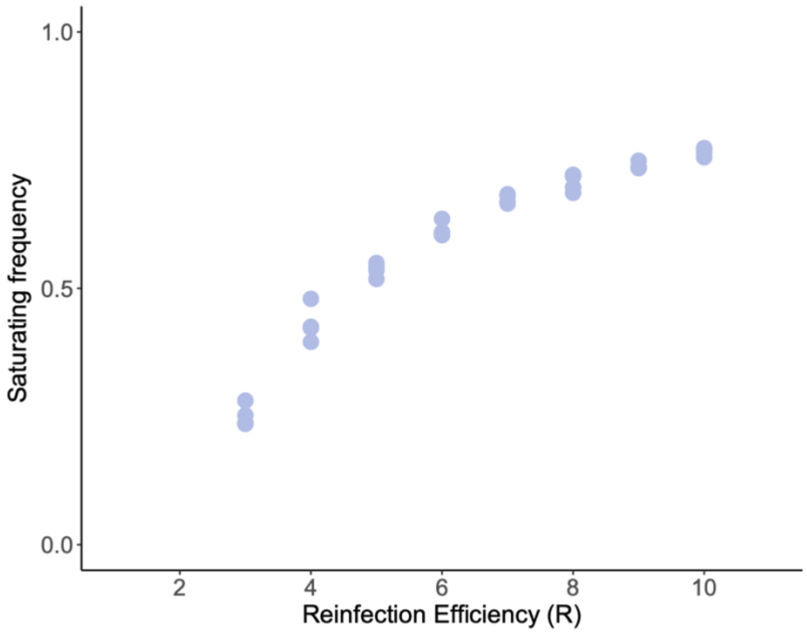

**Figure S11 A phage-plasmid in *Phaeobacter gallaeciensis* DSM26640 and *Phaeobacter piscinae* P18 also shares a highly homologous plasmid backbone but with very different phage structural genes.** These two strains are isolated in geographically close locations but are phylogenetically distinct (different species, average nucleotide identity = 92%).

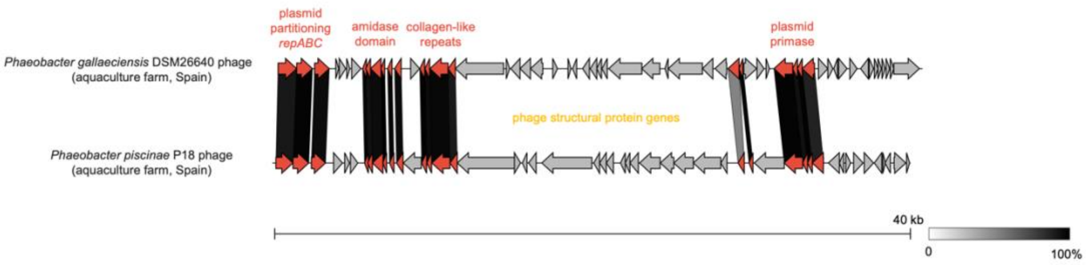

**Figure S12 Geographic distribution of phage repressors that are highly significant hits ($P < 10^{-15}$) of *Tritonibacter mobilis* A3R06 phage-plasmid C1-type repressor gene, identified from currently available environmental metagenomic datasets.**

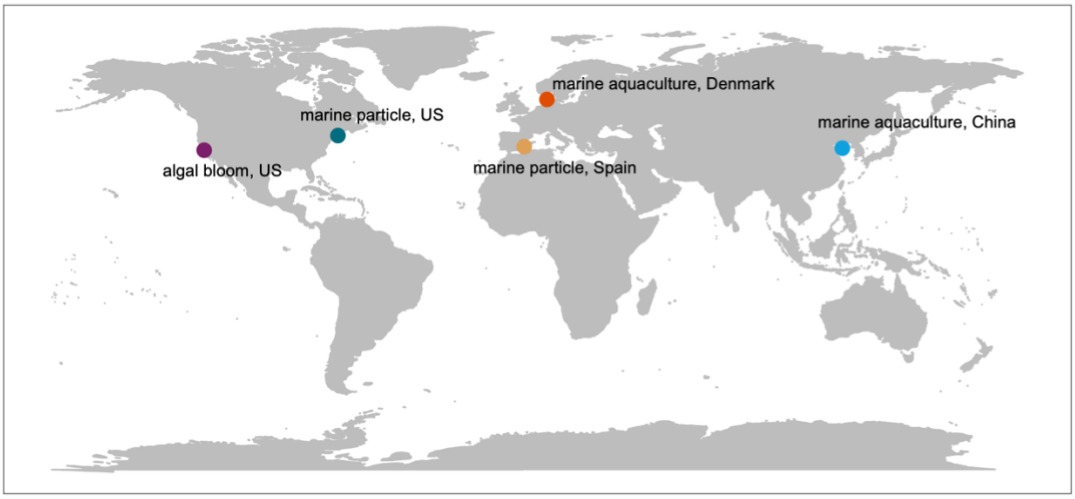

**Figure S13 Differential expression profile for other genes in *Tritonibacter mobilis* A3R06 genome.** Panel 1~4 shows the chromosome, plasmid 1, plasmid 2 and plasmid 3, respectively. Each gene was represented by a bar and is ranked horizontally based on synteny in the genome. Note that each panel has very different scale for the horizontal axis, since there are very different number of genes on each chromosome/plasmid. The gene clusters that are highly up-regulated are related to (A) iron transport, (D) siderophore biosynthesis and (E) fatty acid biosynthesis while those highly down-regulated are related to (B) NADH-ubiquinone oxidoreductase and (C) NADH-ubiquinone oxidoreductase. These changes suggested iron-limitation, oxygen-depletion and modified fatty acid synthesis in a biofilm-type lifestyle in clumpy aggregates. For details of these genes, see Table S2.

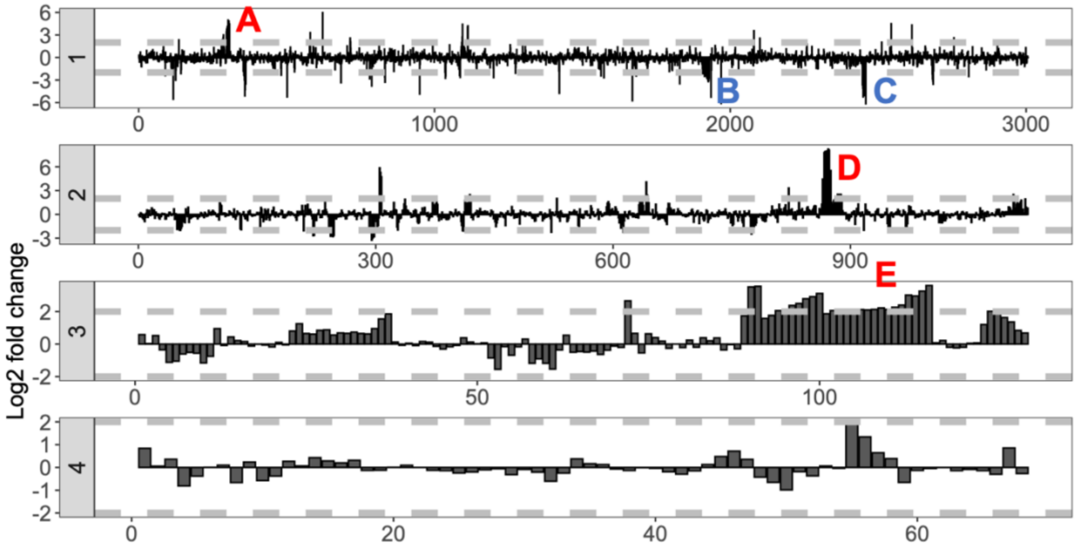
